# DysRegNet: Patient-specific and confounder-aware dysregulated network inference

**DOI:** 10.1101/2022.04.29.490015

**Authors:** Johannes Kersting, Olga Lazareva, Zakaria Louadi, Jan Baumbach, David B. Blumenthal, Markus List

**Affiliations:** Data Science in Systems Biology, TUM School of Life Sciences, Technical University of Munich, Freising, Germany; Chair of Experimental Bioinformatics, TUM School of Life Sciences, Technical University of Munich, Freising, Germany; European Molecular Biology Laboratory, Genome Biology Unit, Heidelberg, Germany; Division of Computational Genomics and Systems Genetics, German Cancer Research Center (DKFZ), Heidelberg, Germany; Wellcome Sanger Institute, Wellcome Genome Campus, Hinxton, UK; Junior Clinical Cooperation Unit Multiparametric methods for early detection of prostate cancer, German Cancer Research Center (DKFZ), Heidelberg, Germany; Institute for Computational Systems Biology, University of Hamburg, Hamburg, Germany; Department of Mathematics and Computer Science, University of Southern Denmark, Odense, Denmark; Department Artificial Intelligence in Biomedical Engineering (AIBE), Friedrich-Alexander-Universität Erlangen-Nürnberg (FAU), Erlangen, Germany

**Author notes:** These authors contributed equally (Johannes Kersting, Olga Lazareva, Zakaria Louadi).

## Abstract

Gene regulation is frequently altered in diseases in unique and patient-specific ways. Hence, personalized strategies have been proposed to infer patient-specific gene-regulatory networks. However, existing methods do not scale well as they often require recomputing the entire network per sample. Moreover, they do not account for clinically important confounding factors such as age, sex, or treatment history. Finally, a user-friendly implementation for the analysis and interpretation of such networks is missing.

We present DysRegNet, a method for inferring patient-specific regulatory alterations (dysregulations) from bulk gene expression profiles. We compared DysRegNet to SSN, a well-known sample-specific network approach. We demonstrate that both SSN and DysRegNet produce interpretable and biologically meaningful networks across various cancer types. In contrast to SSN, DysRegNet can scale to arbitrary sample numbers and highlights the importance of confounders in network inference, revealing an age-specific bias in gene regulation in breast cancer. DysRegNet is available as a Python package (https://github.com/biomedbigdata/DysRegNet_package), and analysis results for eleven TCGA cancer types are available through an interactive web interface (https://exbio.wzw.tum.de/dysregnet).

## Introduction

Gene regulatory network (GRN) inference methods model regulatory relationships based on gene co-expression measures such as (conditional) mutual information or (partial) correlation [1]. A directed network is typically created by limiting the inference to transcription factors (TFs) and their putative target genes. While methods such as GENIE3 [2] or ARACNE [3] identify static GRNs from gene expression data, dynamic methods compare the co-expression in different conditions [4].

Differential expression and co-expression analysis methods designed to compare two groups or more (e.g., disease and control) can typically not account for disease heterogeneity, identify disease subgroups, or describe patient-specific dysregulation patterns. In contrast, methods identifying patientspecific gene expression aberrations in a one-against-all comparison can report sample-specific outlier genes [5, 6]. However, these approaches cannot pinpoint the source of the dysregulation. For instance, a mutated TF may not change in expression but can still behave differently in regulating its target genes, highlighting that co-expression should be considered at the single-patient level.

A few methods for studying patient-specific regulatory patterns have been proposed [7–12]. Most methods calculate the Pearson correlation between two genes before and after adding/removing one sample. Some, such as SSN [10], evaluate the significance of this difference using transformations to z-scores or p-values. P-SSN [11] expands upon this technique by incorporating partial correlation to account for indirect interactions. LIONESS [9] can be adapted to any network inference approach returning a weighted adjacency matrix but does not offer any significance assessment. SWEET [12] extends the LIONESS approach by incorporating a sample-to-sample correlation weight to account for variations in subpopulation sizes but is limited to Pearson correlation as a network inference strategy. Nakazawa et al. [13] define an edge contribution value to extract sub-networks from Bayesian networks inferred from all samples and use this approach successfully for cancer subtyping. We note that existing approaches can not correct for confounders such as sex, age, and origin of the sample, which can impact the analysis at a single sample level. Moreover, the leave-one-sample-out approach is computationally expensive, especially for large cohorts.

These limitations motivated us to develop DysRegNet, a method that first infers linear models from control samples, where the TF expression is considered the explanatory variable and the expression of its target gene the response variable. Subsequently, we consider the residual for each patient sample to determine if the co-expression pattern deviates from the expected value. The linear model allows DysRegNet to correct for known covariates and to compute results (including significance) considerably faster than competing methods (Supplementary Note 1). We show that DysRegNet can infer biologically meaningful patient-specific networks and compare them to results from SSN, a state-of-the-art representative of correlation-based methods.

## Results

### A. Overview of the method

DysRegNet requires a reference GRN and expression data of two groups as input (Figure 1A).

**Fig. 1.**
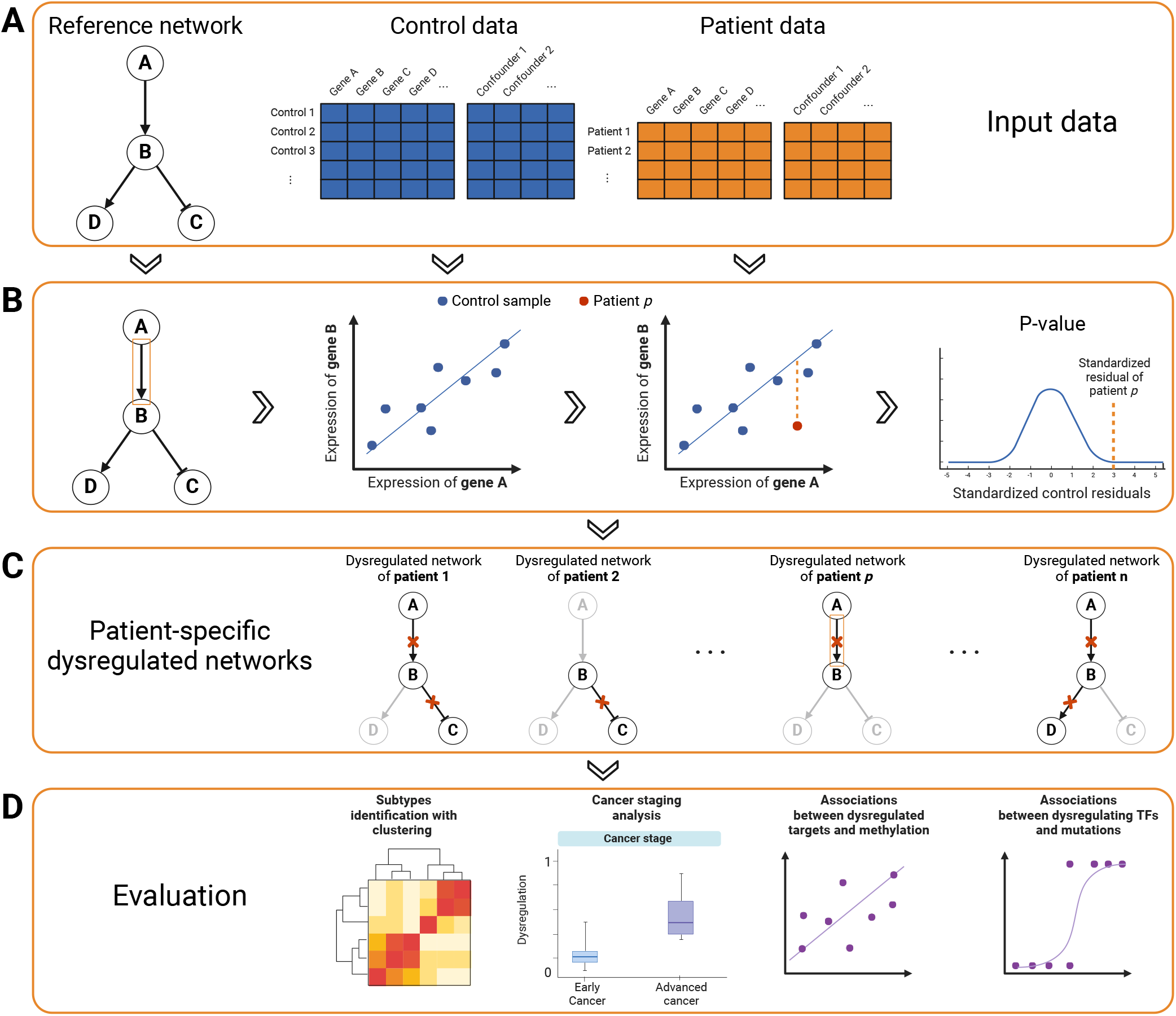
Overview of the method. (A) DysRegNet requires a reference network and expression data for control and patient samples as input. Additionally, multiple sample-level confounders can be provided. For illustration purposes, we assume the reference network has only three edges, with two activating and one repressing interaction. (B) Our method uses the control samples to infer a linear regression model for each edge in the reference network (the illustration focuses on the edge between genes A and B), modeling the expression level of the target gene based on the expression level of the TF and additional confounders if provided. Subsequently, we apply the obtained models to the patient samples, computing a residual for each patient sample. Based on the distribution of the control sample residuals, we assign a p-value to each patient residual. A low p-value suggests that the co-expression pattern of the patient significantly diverges from the control model, leading us to classify the edge as dysregulated. (C) Applying the described procedure to all patient samples and all edges in the reference network leads to the final output of DysRegNet: one network for each patient sample comprising all its dysregulated edges. (D) We evaluated the inferred patient-specific dysregulated networks using known cancer subtypes, cancer stage annotations, additional methylation, and mutation data. Created with BioRender.com

The reference network defines feasible interactions and reduces false positives. It can consist of experimentally confirmed (e.g., HTRIdb [14]) or computationally inferred interactions (e.g., using GENIE3 [2] or ARACNE [3]). The expression data have to be partitioned into two groups: patient and control samples. Control samples define healthy co-expression patterns, which are then used to detect dysregulations in the patient samples. Optionally, DysRegNet can use confounders, such as age or sex, to refine the models further.

Our method fits a linear regression model for every edge in the reference network using the control samples (Figure 1B). Specifically, we model the expression level of the TF as an explanatory variable to estimate the expression level of the target gene. Confounding factors serve as additional explanatory variables. After estimating the model parameters using Ordinary Least Squares, we compute the residual for every patient sample, i.e., the difference between the predicted expression level of the target gene and the observed one. The residual is transformed into a z-score using the distribution of the control sample residuals for standardization. This technique is comparable with an outlier detection task in regression analysis. After evaluating all patients, z-scores are transformed into p-values and corrected for multiple testing (see Methods section J).

The output of our method is a list of predicted dysregulated edges for every patient, which can be integrated into a network with one or several connected components (Figure 1C). It is important to note that previous studies used the term “patient-specific (or sample-specific) regulatory network”. We prefer to call it a patient-specific dysregulated network since we can only identify outliers w.r.t. the original GRN but not learn new edges or a gain of function specific to one sample.

DysRegNet is available as a Python package (https://github.com/biomedbigdata/DysRegNet_package), and analysis results for eleven TCGA cancer types can be interactively explored through a web interface (https://exbio.wzw.tum.de/dysregnet).

### B. Pan-cancer analysis to assess biological relevance

We evaluated DysRegNet using eleven cancer types available in TCGA (see Methods section I) and compared the results to those obtained using SSN (see Discussion section F for a justification for choosing SSN as the reference method in our benchmark). We carried out four analysis types: patient clustering based on the computed networks, promoter methylation of dysregulated targets, dysregulation of mutated TFs, and progression of dysregulation in different cancer stages (Figure 1D).

We used four types of reference networks for the evaluation: GENIE3 individual, GENIE3 shared, experimentally verified interactions from HTRIdb (referred to as *experimental*), and the STRING [15] network. The GENIE3 networks were computationally generated using the GENIE3 method, where the individual network was inferred separately for each cancer type based on its specific control samples, and the shared network comprises the edge intersection of all individual networks. Thus, the shared network is the same for all analyses performed, while the individual networks are cancer-type-specific. The STRING network is not limited to gene-regulatory interactions but considers protein-protein interactions (PPIs). We included a PPI network in the evaluation for a fair comparison with the SSN method that was evaluated using this network. We ran DysRegNet with three additional sample-level confounders, namely sex, age, and ethnicity, which were all previously shown to be strong confounders in certain types of cancers [16–18].

As outlined below, all four analyses suggest that the networks produced by DysRegNet capture biologically relevant signals and, in many cases, to a greater extent than those produced by SSN.

#### B.1. atient-specific networks preserve characterizing features of the cancer type

Even though our method infers an individual network for each patient, aiming to capture its potentially unique dysregulations, we expect similarities between patients of the same cancer type. We assume that patients with the same cancer type have similar dysregulated edges while patients with different cancers have fewer dysregulated edges in common. To investigate this, we evaluated the similarity between all individual patient networks across all cancer types by computing a pairwise overlap coefficient for their edge sets (see Methods section L). We then used spectral clustering to cluster the patients based on their similarity and assigned a label (i.e., the cancer type) to each cluster using the Hungarian algorithm, which maximizes the F1 score of the label-cluster mapping [19]. The evaluation was conducted using the shared GENIE3, HTRIdb, and STRING networks. The individual (cancer-specific) GENIE3 networks were excluded due to high tissue specificity, making conclusions regarding the cancer specificity of the dysregulated networks impossible.

The final F1 scores for DysRegNet and SSN are shown in Figure 2. DysRegNet consistently performed better than SSN in this analysis, particularly when combined with the GENIE3 shared and experimental reference networks. This provides evidence that networks derived from DysRegNet can better capture cancer-type-specific characteristics.

**Fig. 2.**
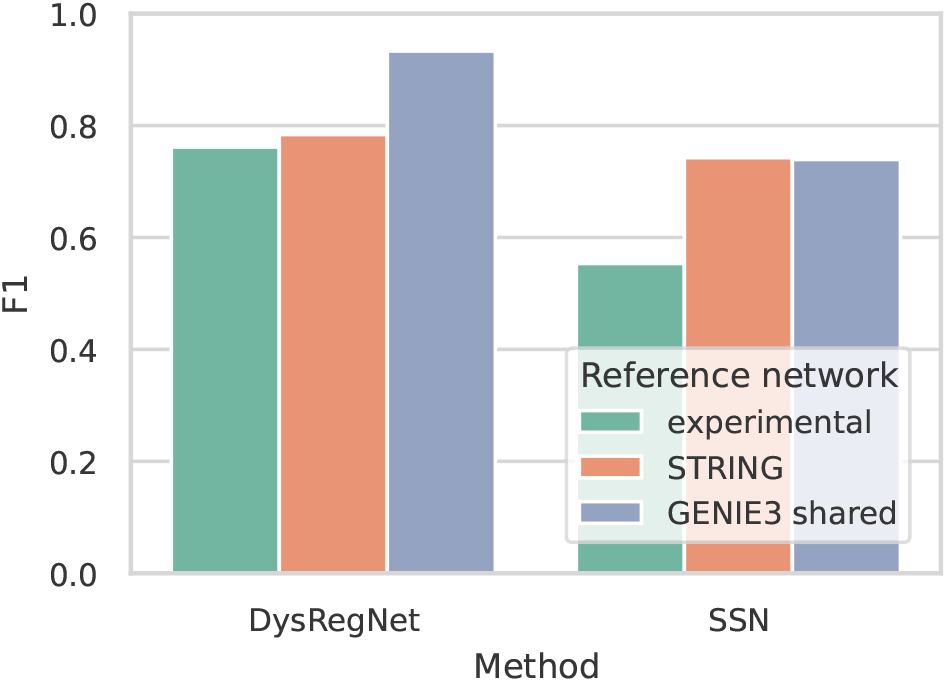
F1 scores assessing the agreement between the patients’ cancer types and their assigned clusters. Higher is better.

#### B.2. Target genes with methylated promoters are more likely to be dysregulated

DNA methylation plays a crucial role in controlling gene expression. For example, when CpG sites undergo methylation within the promoter region, it leads to repressed gene expression because TFs can no longer bind [20]. Therefore, gene promoter methylation is also a diagnostic and prognostic cancer biomarker [21]. Even though changes in promoter methylation represent only one possible cause of dysregulation, we hypothesize that promoter methylation should be correlated with the dysregulation of a target gene across many samples.

Based on this hypothesis, we benchmarked SSN and DysRegNet with respect to the correlation of promoter methylation and patient-specific dysregulation. More specifically, we used linear regression to model the promoter methylation of target genes based on their overall dysregulation. To quantify this overall dysregulation of a target gene in a patient, we defined a dysregulation score as the number of its incoming edges in the patient-specific dysregulated network divided by the number of its incoming edges in the reference network (see Methods section M.1). The dysregulation score, therefore, represents the proportion of TFs associated with a target gene that lost their ability to regulate it. Consequently, a high dysregulation score indicates that TFs generally lost the ability to regulate the target, which is what we would expect in cases where they can no longer bind to the promoter due to methylation. Note that while this would not affect TFs binding to enhancer regions, it suffices to show that dysregulation is related to changes in DNA methylation.

We built two different types of models: a local and a global one. While the local model tests every target gene individually, the global model tries to capture a trend across all target genes by including all of them in one mixed-effect model, adding the target gene as a random intercept (for details, see Methods sections M.2, M.3). An illustration of the differences between the two models can be found in the Supplementary Material Figure S1.

Up to 20% of the tested target genes showed a significant link between their promoter methylation and dysregulation score (Figure 3) in both SSN and DysRegNet. The difference between both methods is relatively small. The results are further corroborated by global significance tests, revealing consistent patterns across all datasets except THCA, LIHC, and KIRC, irrespective of the method or reference network used.

**Fig. 3.**
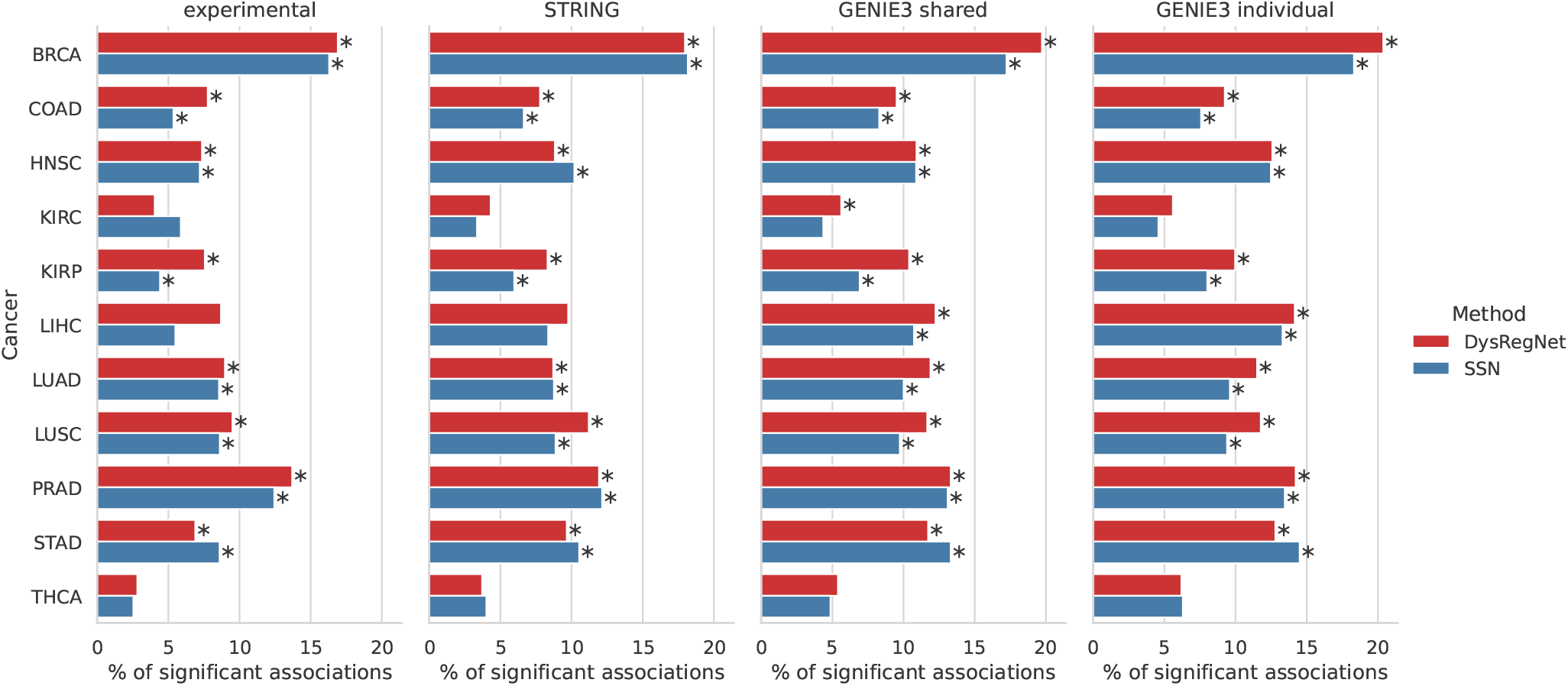
Percentages of target genes with a significant association (Benjamini-Hochberg adjusted p-value ≤ 0.05) between their promoter methylation and target gene dysregulation based on the local models. Bars with a star indicate that the global model (including all target genes) showed a significant association between promoter methylation and target dysregulation.

#### B.3. Mutated TFs have more dysregulated targets

Mutations in TFs are associated with different types of cancer, like lung cancer [22], prostate cancer [23], breast cancer [24], and many others [25]. We hypothesize that some mutated TFs may no longer be able to regulate their target genes, e.g., because they lost the ability to bind their motif.

As a measure, we defined a dysregulation score for TFs analogously to the dysregulation score for target genes used in the methylation analysis, the only difference being that we now focus on outgoing instead of incoming edges. Thus, a high TF dysregulation score indicates that the TF lost the ability to regulate many of its target genes, as expected for some mutated TFs. For this analysis, we modeled the mutation state of a TF based on its dysregulation score. Also, analogously to the methylation analysis, we performed tests using a local and a global model. However, since we treated the mutation state of a TF as a binary outcome, we switched from linear to logistic regression.

Figure 4 shows the results of the local- and global-scale analyses for all cancer types and reference network combinations investigated. The percentages of significant local models exhibit considerably lower and more variable values than those observed in the methylation tests. Nonetheless, the global tests reveal a correlation between mutated TFs and their dysregulation score across numerous scenarios. Again, DysRegNet and SSN perform similarly.

**Fig. 4.**
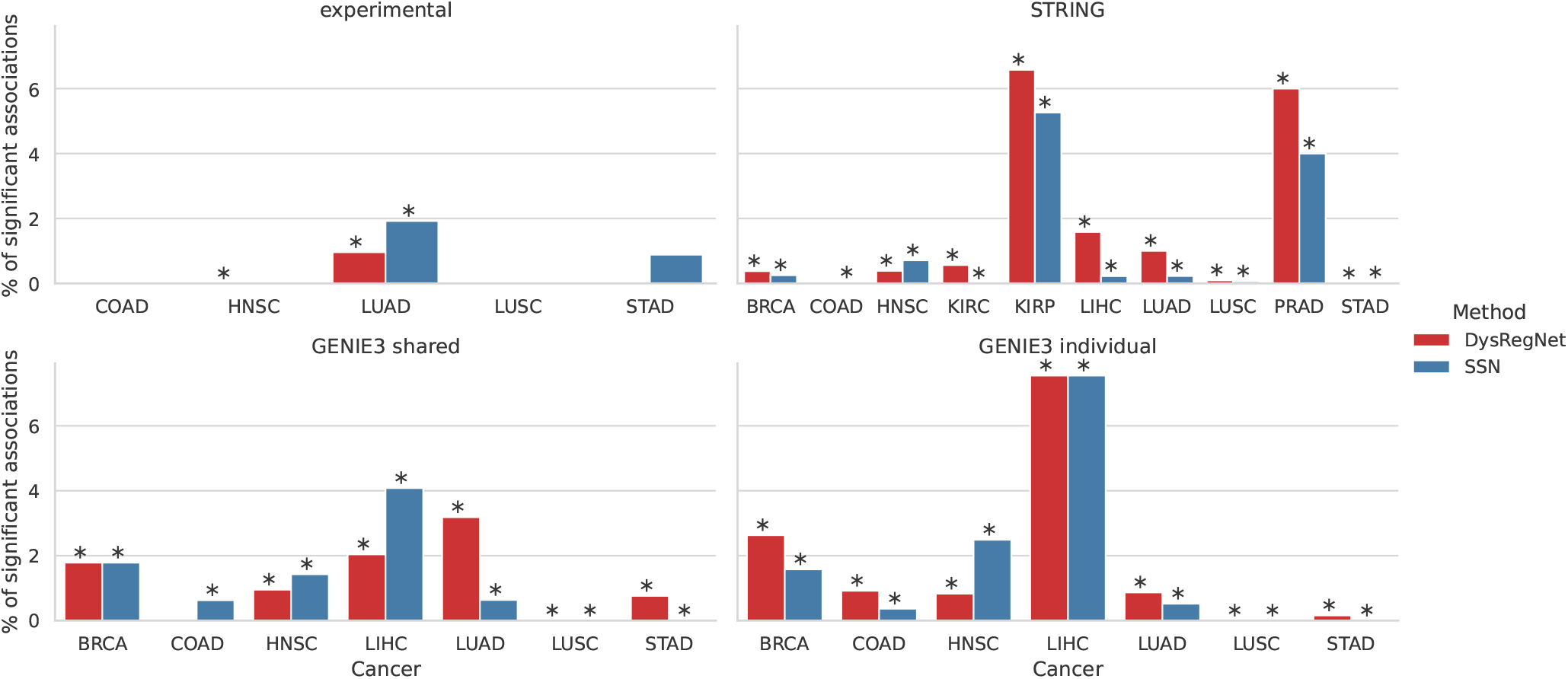
Percentages of TFs with a significant association (Benjamini-Hochberg adjusted p-value ≤ 0.05) between their mutation status and dysregulation. Bars with a star indicate a significant relationship based on the global model. Only cancer types with at least 30 TFs mutated in at least six patients were investigated.

There are multiple factors explaining the disparity in the number of significant tests compared to the methylation analysis: A TF mutation is only one possible source of dysregulation, and only some specific mutations will affect the regulatory ability of TFs. Furthermore, most TFs were only mutated in a handful of patients, reducing the statistical power of our analyses.

Because of this, we still see the significant relationship between mutated TFs and their dysregulation indicated by global tests as further evidence of the biological meaningfulness of patient-specific networks.

#### B.4. Patients with advanced stages of cancer exhibit increased dysregulation

Due to the accumulation of mutations [26] and progressing epigenetic reprogramming of gene regulation in cancer [27], we expect to observe increased gene dysregulation in the late stages of cancer compared to the early stages.

To investigate this assumption, we divided patients into early-stage and late-stage groups for each cancer type (see Methods section M.6). We then applied a one-sided Mann-Whitney U test to asses whether the dysregulation scores of TFs would be increased in the late-stage groups.

Except for COAD and BRCA, large percentages of TFs showed increased dysregulated in late-stage patients, exceeding 80% for LIHC Figure 5. Compared to SSN, DysRegNetinferred networks generally contained more TFs with increased dysregulation in the advanced stages.

**Fig. 5.**
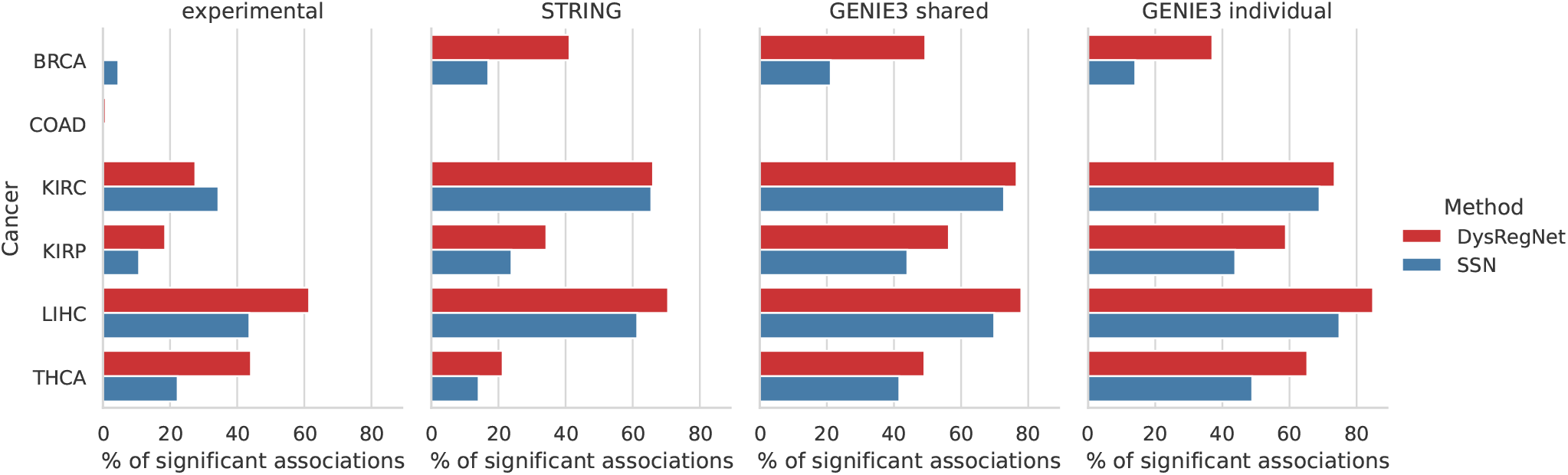
Percentages of TFs with significantly (Benjamini-Hochberg adjusted p-value ≤ 0.05) increased dysregulation in advanced cancer stages. Only cancer types with at least 30 patients in both (early-stage and late-stage) groups were tested.

#### B.5. Confounders significantly impact model inference

A distinguishing feature of DysRegNet is the ability to correct for sample-level confounders. To assess if including a patient’s sex, age, and ethnicity has an impact, we assessed the covariates model coefficients Figure 6. Unsurprisingly, the expression of the TF produced the most significant p-values. However, the confounders noticeably impacted the regression models in various cancer types, such as age in the breast cancer (BRCA) dataset, sex in liver hepatocellular carcinoma (LIHC), and ethnicity in stomach adenocarcinoma (STAD) and lung adenocarcinoma (LUAD). In BRCA, *NOX4*, the target gene most strongly positively associated with age, was indeed observed in previous studies to increase with age [28– 30]. In LIHC, *KDM5C* was the target gene associated with the most significant p-value for sex. *KDM5C* is encoded on the X chromosome and is known to have higher expression in females than in males [31, 32], which was again reflected by the sign of the estimated coefficient. This indicates that patient-level confounding factors influence the expression data, which DysRegNet can efficiently incorporate into its model construction and patient-specific network inference process.

**Fig. 6.**
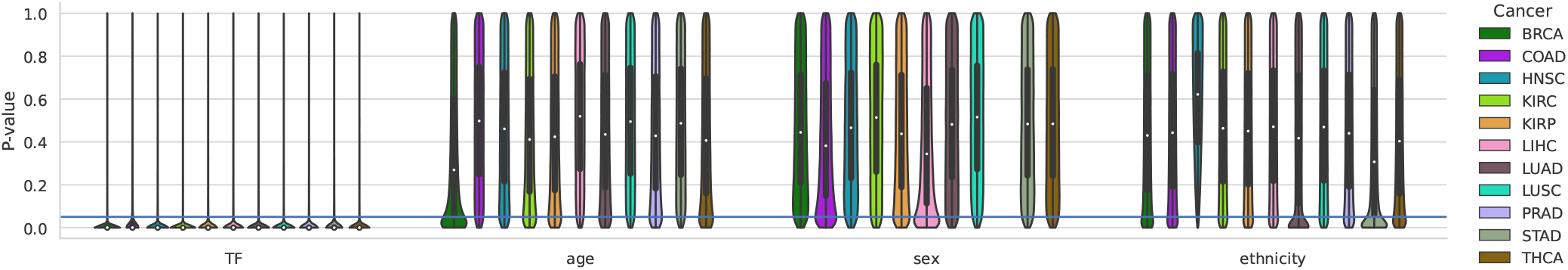
Coefficient p-value distributions for different cancer types. DysRegNet results are based on the GENIE3 shared reference network. A single violin summarizes the p-values of a selected coefficient and cancer type across all linear models (one for each edge in the reference network). A confounding variable lacking meaningful predictive power will yield a uniform distribution of p-values. Conversely, influential confounders are expected to generate a disproportionate number of small p-values, manifesting as a pronounced concentration at the lower end of the p-value distribution, akin to a “belly” in the violin plot. Note that there are no p-values for sex in prostate adenocarcinoma (PRAD) as it inherently includes only male samples.

## Discussion

### C. Validity of model assumptions

Since DysRegNet is an outlier detection approach, it will be especially susceptible to technical biases. Furthermore, users should remember that the method is built based on the following assumptions: (1) the target gene expression can be predicted based on TF expression, and (2) the residuals of the linear model follow a normal distribution. The first assumption typically does not hold for PPI networks. It will also not hold for all TF-target gene pairs in a GRN. The activity of a TF can be influenced by many factors, such as interactions with other proteins, chromatin accessibility, or post-translational modifications, which are not reflected in expression data [33, 34]. We thus recommend evaluating the goodness of fit of the underlying regression model before considering an edge as potentially dysregulated. Our Python package directly supports this by setting an *R*^2^ cut-off. Similarly, we implemented a test to evaluate whether the residuals follow a normal distribution [35, 36]. This allows users to focus only on regulatory interactions that adhere to our second assumption.

### D. Importance of adjusting for confounders

Our results suggest that DysRegNet can account for typical confounders such as age and sex. However, it should be noted that these confounders do not seem to impact all data sets to the same extent. Age, for example, only affected the BRCA dataset. This finding is well-aligned with a recent study on the effect of demographic confounders on cohort-level network inference tools where age was identified as a strong confounder on network inference in BRCA based on data from two independent cohorts (TCGA and METABRIC)[37]. Given that BRCA has notably more control samples (113) than other cancer types (see Table S1), we assume that certain effects require a sufficiently large sample to be detected. The same principle applies to categorical confounders such as sex or ethnicity, where aside from the overall sample size, each category must appear frequently enough to accurately estimate its effect.

### E. Influence of different reference networks

The initial reference network is a necessary input for DysRegNet. Our analysis considered three types: computationally inferred regulatory networks from GENIE3, an experimentally validated network from HTRIdb, and a PPI network from STRING. We compared these networks in four different contexts: patient clustering (i), promoter methylation of dysregulated target genes (ii), dysregulation of mutated TFs (iii), and progression of dysregulation in different cancer stages (iv).

The choice of the network depends on the research question. Users can expect computationally inferred networks to favor false positives, while experimentally validated networks may include more false negatives. In our analyses, the choice of the reference network seldom changed overall patterns. Our analyses produced better results when DysRegNet and SSN were combined with the computationally inferred GENIE3 networks, followed by the STRING and the experimentally validated network. A possible explanation is that the GENIE3 networks were constructed based on the same expression data used for the subsequent patient-specific network inference. Consequently, they more accurately reflect co-expression patterns observed in the data. As shown in Figure S2, the *R*^2^ values obtained by the linear regression models of DysRegNet were generally higher with the computationally inferred networks, indicating a better model fit.

Gene regulation differs across tissues and cell types [38]. Generic GRNs such as STRING and experimentally validated networks likely include regulatory interactions that may not be relevant to the tissues and datasets under study. In addition, these networks might encompass valid yet highly complex or non-linear relationships, which are not easily captured by techniques such as linear regression or Pearson correlation. Both scenarios can lead to edges that cannot be reliably modeled with DysRegNet or the available data, potentially impacting the meaningfulness of the results. Because of this, the DysRegNet Python package provides an option to only consider edges, which can be modeled with a certain *R*^2^ value or higher.

Furthermore, it is important to note that the STRING network violates some of our modeling assumptions. For DysRegNet, we expect a gene regulatory reference network of directed TF-target gene relationships. However, the edges in the STRING network are neither directed nor are they actual TF-target gene pairs. While the overall co-expression modeling approach may still be valid, as we expect co-expression patterns between interacting proteins, our downstream measurements, such as dysregulation scores for TFs and target genes, lose interpretability.

### F. Evaluation

We evaluated our approach against the original single-sample GRN inference method SSN. Although we compared our method only to this approach, numerous other methods adopted a comparable procedure. Lee et al. [8] extended the SSN approach by integrating multi-omics data such as copy number variation or DNA methylation. Due to the high similarity of methods and lack of public source code, a circumstance shared with Nakazawa et al. [13], we did not include those methods in the evaluation. Furthermore, the available code of P-SSN [11] requires a special reference network with a list of potentially indirectly interacting genes for each edge. Since P-SSN does not use standard regulatory networks as input, a proper comparison with DysRegNet was impossible. While DysRegNet identifies dysregulation compared to a healthy background, LIONESS [9] and SWEET [12] do not utilize a dedicated set of control samples but capture truly sample-specific interactions. The right tool choice thus depends on the question and the availability of control samples, which are often not available at a sufficient number (at least 20 samples are needed for a robust co-expression analysis [39]). For studies without controls, utilizing independent tissue-specific expression data of healthy individuals can be an alternative. Park et al. [7], for instance, employ a methodology closely related to the SSN approach where they leverage GTEx [40] data. However, cross-study batch effects need to be accounted for in this case.

Despite recent attempts to systematically compare various sample-specific network inference methods [12, 41], a proper evaluation remains a major challenge. We considered a variety of cancer expression datasets and networks across different complementary scenarios. Overall, our results indicate that DysRegNet and SSN produce biologically meaningful results. However, our evaluation is limited by the absence of ground truth and discrepancies in interpreting the networks generated by different methods [42]. For example, we expect a higher percentage of dysregulated TFs in advanced cancer stages, while the true fraction or upper limit is unknown. Similarly, SSN networks generally comprise more dysregulated edges than those of DysRegNet, even though we used the same p-value cut-off and multiple testing correction procedures for both methods (Figure S3). While this could suggest that DysRegNet generates fewer false positives, we can not prove this without knowing the ground truth.

### G. Outlook

A promising direction for further research is a more in-depth investigation of the connection between dysregulations and mutations. This study investigated whether mutated TFs will be more frequently involved in dysregulation. However, unlike in the promoter methylation analysis, the association between dysregulation and mutations was less visible. We found significant associations for 7% of the tested TFs at most. Our analysis did not consider that not every somatic mutation will affect the regulatory function of a TF. A more specific analysis could focus on known or patientspecific mutations within the DNA binding domains of TFs in the promoter or enhancer regions of target genes and model their impact on expression changes [43]. Furthermore, somatic mutations in cancer frequently affect the splice site and can cause isoform switches [44, 45]. Our analysis was performed at the gene level, but a deeper analysis at the isoform or transcript level would help explain a larger fraction of the identified dysregulated edges.

Another interesting application of the method is in studying rare and undiagnosed diseases, where the focus is often on the unique differences of a single sample. The current rate of genetically diagnosed rare disorders is approximately 25 to 50% [5]. Thus, DysRegNet provides a novel opportunity to expand our knowledge of such disorders.

Single-patient network extraction is a promising research direction toward precision medicine as it highlights potential treatment targets in a personalized fashion. A plethora of computational methods have been developed for drug target identification and drug repurposing [46]. Many of these methods already use networks as input [47–49] and embrace concepts of network pharmacology [50]. Typically, such methods extract disease modules after connecting patient- or disease-specific seed genes. Currently, seed gene selection is typically based on literature or patient-specific mutation or expression profiles. To our knowledge, patient-specific dysregulation and co-expression analysis have thus far not been employed to identify seed genes, suggesting a promising direction for further research.

### H. Conclusion

Aberrant TF regulation is an important mechanism in complex diseases such as cancer. Rather than focusing on the aberrant expression of TFs or their target genes, it is worthwhile to study which specific interactions of a TF are affected to gain a more detailed insight into the underlying pathomechanisms. Many molecular changes can lead to the same outcome; hence, it is vital to study dysregulation in a patient- or sample-specific manner. With DysRegNet, we present a novel approach that delineates individual TF-regulatory changes in relation to a control cohort. In contrast to competing methods, DysRegNet uses linear models to account for confounders and residual-derived z-scores to assess significance. Due to the latter, DysRegNet scales efficiently to an arbitrary number of samples. We have shown that DysRegNet results are robust across template networks and produce meaningful insights into cancer biology. DysRegNet may hence serve as a precision medicine tool for identifying drug targets in oncology and beyond.

## Materials and Methods

### I. Data preprocessing

All data from The Cancer Genome Atlas Program (TCGA) (comprising gene expression profiles based on bulk RNA-seq, methylation, mutation, and sample metadata) were acquired from the XENA browser (https://xena.ucsc.edu/) [51]. We included eleven cancers with at least 30 control samples available (labeled “Solid Tissue Normal”). We used log_2_(*tpm* + 0.001) normalized counts of the PANCAN cohort as a gene expression dataset (see Table S1 for the exact sample numbers). We retained only genes expressed in at least 80 % of the patients of a cancer type. Subsequently, we standardized the expression data gene-wise based on the mean and standard deviation of the cancer-specific control samples.

The three sample-level confounders used for DysRegNet (sex, age, and ethnicity) were obtained from the curated clinical data. Missing values for age were replaced with the mean age across all samples of a cancer type.

Illumina 450k DNA methylation array data was filtered for CpGs associated with promoter regions (according to Illumina’s annotation). We then calculated the mean of all methylation *β*-values associated with the same gene to obtain a methylation value for each gene in each sample.

Somatic mutations were mapped to their genes. In our analysis, we considered mutations of 12 different effect categories: Missense, nonsense, intron, frameshift deletion, frameshift insertion, splice site, in-frame deletion, in-frame insertion, RNA, translation start site, nonstop mutation, and large deletion.

### J. The statistical model behind DysRegNet

We define a reference network *N* = (*G, T, E*), where *G* is a set of genes, *T* ⊆ *G* is a set of TFs, and *E* ⊆ *T* × *G* is a set of edges connecting TFs *t* ∈ *T* to target genes *g* ∈ *G*. The role of the reference network is to limit the search space for potentially dysregulated edges and to provide prior information about expected healthy regulations. We discuss possible choices for the template network in section K.

For every pair of connected nodes (*t*_*i*_, *g*_*j*_), the relationship between the expression profiles of a TF *t*_*i*_ and a target gene *g*_*j*_ can be modeled as:

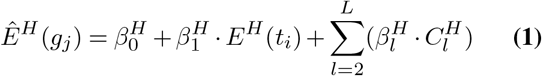

where *E*^*H*^ (*t*_*i*_) is the expression of a TF *t*_*i*_ in a cohort of healthy controls, Ê^*H*^ (*g*_*j*_) is the expected expression of a target gene *g*_*j*_ in a cohort of control samples, 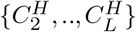 is a set of available covariates such as age, ethnicity, and sex, 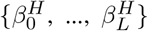 are coefficients estimated with an ordinary least squares model.

An edge *e*_*ij*_ = (*t*^*i*^, *g*^*j*^) is dysregulated for a patient *p* if the edge exists in the reference network *N*, and, for patient *p*, the expression of *g*_*j*_ cannot be reliably estimated using the model from Equation 1. Formally, this means that the expected expression of 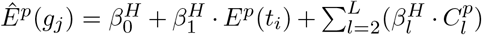 is *significantly* different from the actual value of *E*^*p*^(*g*_*j*_). This difference can be defined as a residual of the model, i.e.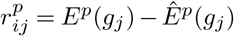, which can be converted to a z-score using the following transformation:

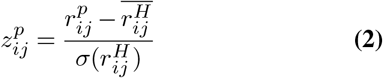

where 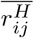 and 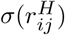 are the mean and standard deviation of the control cohort residuals 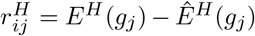 . The z-scores are converted to p-values using the standard normal distribution and subsequently corrected for multiple hypothesis testing regarding the number of patients with the Bonferroni correction procedure at a 0.01 significance level.

### K. Reference networks

In our analyses, we used three types of reference networks: computationally inferred networks using GENIE3 [2], an experimentally derived regulatory network from the Human Transcriptional Regulation Interactions database (HTRIdb) [14], and a PPI network from STRING [15].

#### GENIE3

GENIE3 uses an ensemble of trees to estimate the strength of the regulatory relationship between all possible TF-target gene pairs. A list of 1639 human TFs was used from Lambert et al. [52] (http://humantfs.ccbr.utoronto.ca/) to limit the search space. To obtain cancer-specific reference networks, we ran the GENIE3 R package (https://github.com/aertslab/GENIE3, version 1.20.0) individually on the control samples of each cancer type. The shared network was derived by summing up the edge importance scores of all cancer-specific networks. For all GENIE3-inferred networks, we retained only the top 100,000 highest-scoring edges. We chose this cut-off to obtain networks that fall in between the relatively small HTRIdb and the much larger STRING network in terms of size.

#### HTRIdb

The Transcriptional Regulation Interactions database (http://www.lbbc.ibb.unesp.br/htri) is an open-access database of experimentally validated TFtarget gene interactions. The database provides information about regulation interactions among 284 TFs and 18,302 target genes detected by 14 distinct techniques [14]. Namely, chromatin immunoprecipitation, concatenate chromatin immunoprecipitation, CpG chromatin immunoprecipitation, DNA affinity chromatography, DNA affinity precipitation assay, DNase I footprinting, electrophoretic mobility shift assay, southwestern blotting, streptavidin chromatin immunoprecipitation, surface plasmon resonance and yeast one-hybrid assay, chromatin immunoprecipitation coupled with microarray (ChIP-chip) or chromatin immunoprecipitation coupled with deep sequencing (ChIP-seq).

#### STRING

The STRING database (http://string-db.org/) is dedicated to protein-protein interactions. It was included in our assessment for a fair comparison between DysRegNet and SSN [10] (see section O), which was originally evaluated using the STRING network. Following the described methodology, we also considered high-confidence interactions with a combined score larger than 0.9, retaining a total of 197,969 edges. The combined score is an aggregate of 7 channels (neighborhood, fusion, co-occurrence, co-expression, experimental, database, text mining).

### L. Patient clustering

We evaluated the similarity between patient-specific networks by computing a pairwise overlap coefficient for the set of dysregulated edges, i.e.:

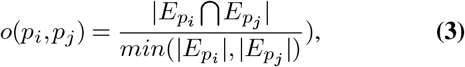

where 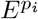 and *Ep*_*j*_ are sets of the dysregulated edges for patients *i* and *j*, respectively. The resulting similarity matrix was used to cluster the patients with spectral clustering implemented in scikit-learn (https://github.com/scikit-learn/scikit-learn, version 1.1.3, [53]. We assigned cancer-type labels to the clusters, maximizing the F1 score using the Hungarian algorithm [19] implemented in the munkres Python package (https://github.com/bmc/munkres, version 1.1.4).

### M. Hypothesis testing for mutation, methylation, and cancer stage analysis

#### M.1. Dysregulation scores

We compute dysregulation scores to quantify the dysregulation of a gene (either a TF or a target gene) in an individual patient. For a target gene *g*, the dysregulation score is defined as 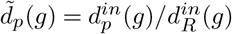 where 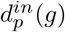 is the in-degree of *g* in the dysregulated network of patient *p* and 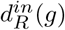 is the in-degree of *g* in the reference network. Analogously, for a transcription factor *t*, the dysregulation score is defined as 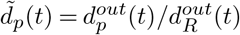 where 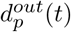is the out-degree of *t* in the dysregulated network of patient *p* and 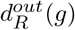 is the out-degree of *t* in the reference network.

#### M.2. DNA methylation local model

To model the relationship between promoter DNA methylation and target gene dysregulation, we used the following linear model:

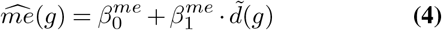

Here, *me*(*g*) = [*me*_1_(*g*),…, *me*_*P*_ (*g*)] is the vector of the average (across CPGs) promoter DNA methylation (see section I) of target gene *g* for all *P* patients and 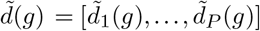 is the vector of *g*’s dysregulation scores. The slope coefficients 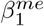 were tested for significance with the null hypothesis 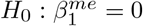 . The p-values were then corrected using the Benjamini-Hochberg method.

The linear models were estimated using the ordinary least squares implementation in the statsmodels Python package (https://github.com/statsmodels/statsmodels, version 0.13.5).

#### M.3. DNA methylation global model

While Equation 4 allows us to test every target gene individually, we also applied a linear mixed-effect model to investigate the global association between promoter methylation and dysregulation across all target genes in the reference network. For this, we built a model with a random intercept coefficient for each target, assuming different baseline methylation levels:

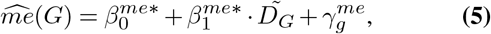

where *me*(*G*) are the average (across CPGs) promoter DNA methylation values for any 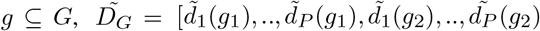 are target gene dysregulation scores across all target genes and patients, and 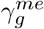 is a random intercept for each target gene. The slope coefficient 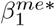 was tested for significance with the null hypothesis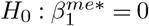.

The linear mixed-effect models were estimated using the implementation in the statsmodels Python package (https://github.com/statsmodels/statsmodels, version 0.13.5).

#### M.4. Mutation local model

We tested the association between a TF’s dysregulation score and its mutation status using a logistic regression model:

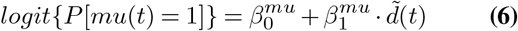

Here, *mu*(*t*) = [*mu*_1_(*t*), …, *mu*_*P*_ (*t*)] is a binary vector indicating the mutation status (1 indicates the presence of at least one mutation, 0 means no mutation) of the TF *t* for all *P* patients and 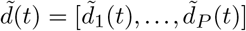 is the vector of *t*’s dysregulation scores.

The slope coefficient 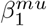 was tested for significance with the null hypothesis 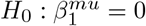 . The p-values were then corrected using the Benjamini-Hochberg method. We only tested TFs that were mutated in at least six patients and only considered cancer types with a least 30 TFs fulfilling this criterion.

The logistic regression models were estimated using the implementation in the statsmodels Python package (https://github.com/statsmodels/statsmodels, version 0.13.5).

#### M.5. Mutation global model

To evaluate the global relationship between TF mutations and dysregulation, we used a logistic mixed-effect model with a random intercept coefficient for each TF, assuming different baseline mutation loads:

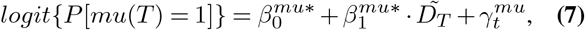

where *mu*(*T*) are binary mutation statuses for any *t* ⊆ *T*, 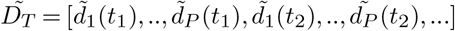 are TF dysregulation scores, and 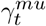 is a random intercept for every TF. 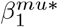 was tested for significance with the null hypothesis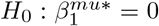. We only tested TFs that were mutated in at least six patients and only considered cancer types with a least 30 TFs fulfilling this criterion.

The logistic mixed-effect models were estimated using the implementation in the pymer4 Python package (https://github.com/ejolly/pymer4, version 0.8.0)

#### M.6. Hypothesis testing for cancer stage analysis

We separated the samples of each cancer into two groups: an early-stage group (stage I) and a late-stage group (stage III or stage IV). For better separability, we excluded stage II samples from this analysis since they pose an intermediate setting that can resemble either stage I or III too closely. We conducted cancer stage analysis exclusively for types of cancer with a minimum of 30 patients in both groups to ensure the preservation of statistical power. We then performed a one-sided Mann-Whitney U test, implemented in scikit-learn (https://github.com/scikit-learn/scikit-learn, version 1.1.3, [53]), for every TF in the reference network to assess whether its dysregulation scores are larger in the late-stage compared to the early-stage group. We corrected the obtained p-values for multiple testing with the Benjamini-Hochberg method.

### N. Coefficient p-values

Two-tailed p-values for the coefficients of DysRegNet’s linear models were obtained based on the coefficient’s t-statistic. They can be interpreted as the probability that the coefficient is zero and thus has no predictive power. Multi-label categorical confounders, such as ethnicity, resulted in multiple (*l* − 1, where *l* is the number of different categories) coefficients/p-values per model, which, for visualization purposes, were summarized in one distribution.

### O. SSN

SSN, described by Liu et al. [10], calculates the correlation difference introduced by adding one case sample to a set of control samples. For each case sample and each pair of genes, the following score is computed:

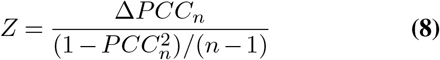

where Δ*PCC*_*n*_ = *PCC*_*n*+1_ − *PCC*_*n*_ is the difference between the Pearson correlation coefficients calculated on the control samples (*PCC*_*n*_) and the control samples with one case sample (*PCC*_*n*+1_), and *n* is the number of control samples. We further converted the *Z* values into p-values with a *Z*-test, as described by Liu et al. Next, we corrected the p-values for multiple hypothesis testing using the Bonferroni method. We set a cut-off that defines a dysregulated edge at a corrected p-value of 0.01 or lower, equal to the cut-off selected for DysRegNet.

### P. Web interface

The dysregulated networks of the eleven studied TCGA cancers can be interactively explored using a web interface (https://exbio.wzw.tum.de/dysregnet), which was built with Plotly Dash (https://plotly.com/dash/, version 2.0.0), the Cytoscape.js [54] wrapper Dash Cytoscape (https://dash.plotly.com/cytoscape, version 0.2.0), Dash Bio (https://dash.plotly.com/dash-bio, version v0.2.0) and a Neo4j database (https://neo4j.com/, version 5.11.0). We inferred the visualized networks using the GENIE3 shared reference network.

Since the underlying network is vast and highly connected, the interface is centered around individually selected query genes. Only the regulatory connections between those genes and their targets or sources are displayed to keep the resulting network compact and tidy. Further query genes can be added to expand the graph in directions of interest.

We display the fraction of patients with a dysregulation for each regulatory connection, which is directly depicted by the corresponding edge in the graph network. This metric can also be compared visually between different cancer types. Furthermore, the web interface incorporates information about the gene mutation frequency and mean promoter DNA methylation. Heatmaps allow the investigation of the DNA methylation status and the significance of a dysregulation on the patient level.

To prevent the underlying graph structure from becoming too large, the maximum number of displayed edges is capped, and edges can be filtered by their fraction of dysregulated patients and their type. In case a user is interested in the full, unfiltered graph, it can be downloaded as a CSV file.

The displayed network can be directly exported to the online systems medicine platform Drugst.One [55] to obtain drugs targeting the dysregulated genes.

### Q. Python package

An implementation of DysRegNet as described in section J is available as an easy-to-access Python package (https://exbio.wzw.tum.de/dysregnet). The linear regression modeling, as well as coefficient p-value and *R*^2^ value calculation, were implemented based on the ordinary least squares implementation in the statsmodels Python package (https://github.com/statsmodels/statsmodels).

Our package also provides some additional features, which we did not use in this study for better comparability with SSN. This includes a goodness of fit filter to ignore edges with a low *R*^2^ value and the normality filter [35, 36] implemented in the scipy Python package (https://github.com/scipy/scipy) to test the assumption of normally distributed control sample residuals. Furthermore, the Python package can distinguish between four possible scenarios of dysregulation shown in Figure S5: suppressed activation (1), amplified activation (2), amplified repression (3), and suppressed repression (4). The Python package can be used to only consider scenarios 1 and 4, which correspond to a reduced response towards the TF expression rather than an amplified one (scenarios 2 and 3).

## Acknowledgments

The results shown here are in whole or part based upon data generated by the TCGA Research Network: https://www.cancer.gov/tcga. Contributions from Z.L, J.B, and M.L are funded by the German Federal Ministry of Education and Research (BMBF) within the framework of the e:Med research and funding concept [grant 01ZX1908A (Sys_CARE)]. Contributions by O.L. are funded by the Bavarian State Ministry of Science and the Arts within the framework coordinated by the Bavarian Research Institute for Digital Transformation (bidt, Doctoral Fellow). The project received funding from the Deutsche Forschungsgemeinschaft (DFG, German Research Foundation) [422216132]. This project has received funding from the European Union under grant agreements No. 101057619 and No 777111. Views and opinions expressed are however those of the author(s) only and do not necessarily reflect those of the European Union or European Health and Digital Executive Agency (HADEA). Neither the European Union nor the granting authority can be held responsible for them. This work was also partly supported by the Swiss State Secretariat for Education, Research and Innovation (SERI) under contract No. 22.00115. Furthermore, this work was supported by the German Federal Ministry of Education and Research (BMBF) within the framework of the e:Med research and funding concept (grant 01ZX1910D). JB was partially funded by his VILLUM Young Investigator Grant nr.13154. D.B.B. was supported by the German Federal Ministry of Education and Research (BMBF) within the framework of the CompLS research and funding concept [grant 031L0309A (NetMap)].

## Data Availability

The datasets and computer code produced in this study are available in the following databases:

- expression, mutation, methylation, and sample metadata: TCGA Pan-Cancer (PANCAN) cohort, XENA browser (https://xenabrowser.net/datapages/)
- PPI network: STRING database (http://string-db.org/)
- Experimentally validated regulatory network: Transcriptional Regulation Interactions database (originally obtained from http://www.lbbc.ibb.unesp.br/htri, currently available at https://rescued.omnipathdb.org/).
- List of human TFs: http://humantfs.ccbr.utoronto.ca/
- Python package: GitHub (https://github.com/biomedbigdata/DysRegNet_package)
- Analysis scripts: GitHub (https://github.com/biomedbigdata/DysRegNet_workflow)

---

**Table S1.** Number of available control samples (additionally separated by sex) and patient samples for each studied cancer type.

| Cancer | Control samples | Female control samples | Male control samples | Patient samples |
| --- | --- | --- | --- | --- |
| BRCA | 113 | 112 | 1 | 1098 |
| COAD | 41 | 21 | 20 | 288 |
| HNSC | 44 | 14 | 30 | 520 |
| KIRC | 72 | 20 | 52 | 531 |
| KIRP | 32 | 10 | 22 | 289 |
| LIHC | 50 | 22 | 28 | 371 |
| LUAD | 59 | 34 | 25 | 515 |
| LUSC | 50 | 14 | 36 | 498 |
| PRAD | 52 | 0 | 52 | 496 |
| STAD | 36 | 13 | 23 | 414 |
| THCA | 59 | 42 | 17 | 512 |

**Fig. S1.**
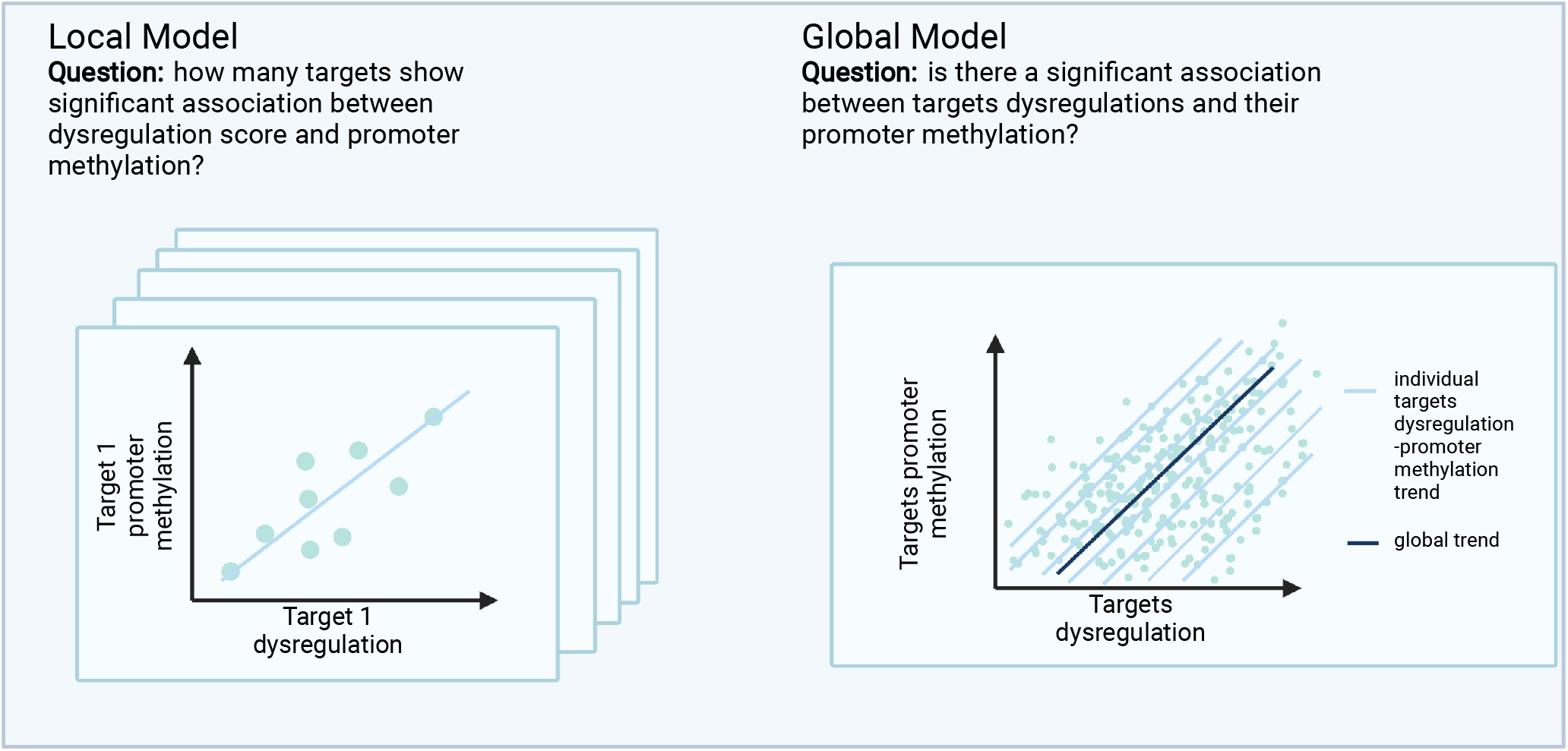
Local and global models for methylation-dysregulation association studies.

**Fig. S2.**
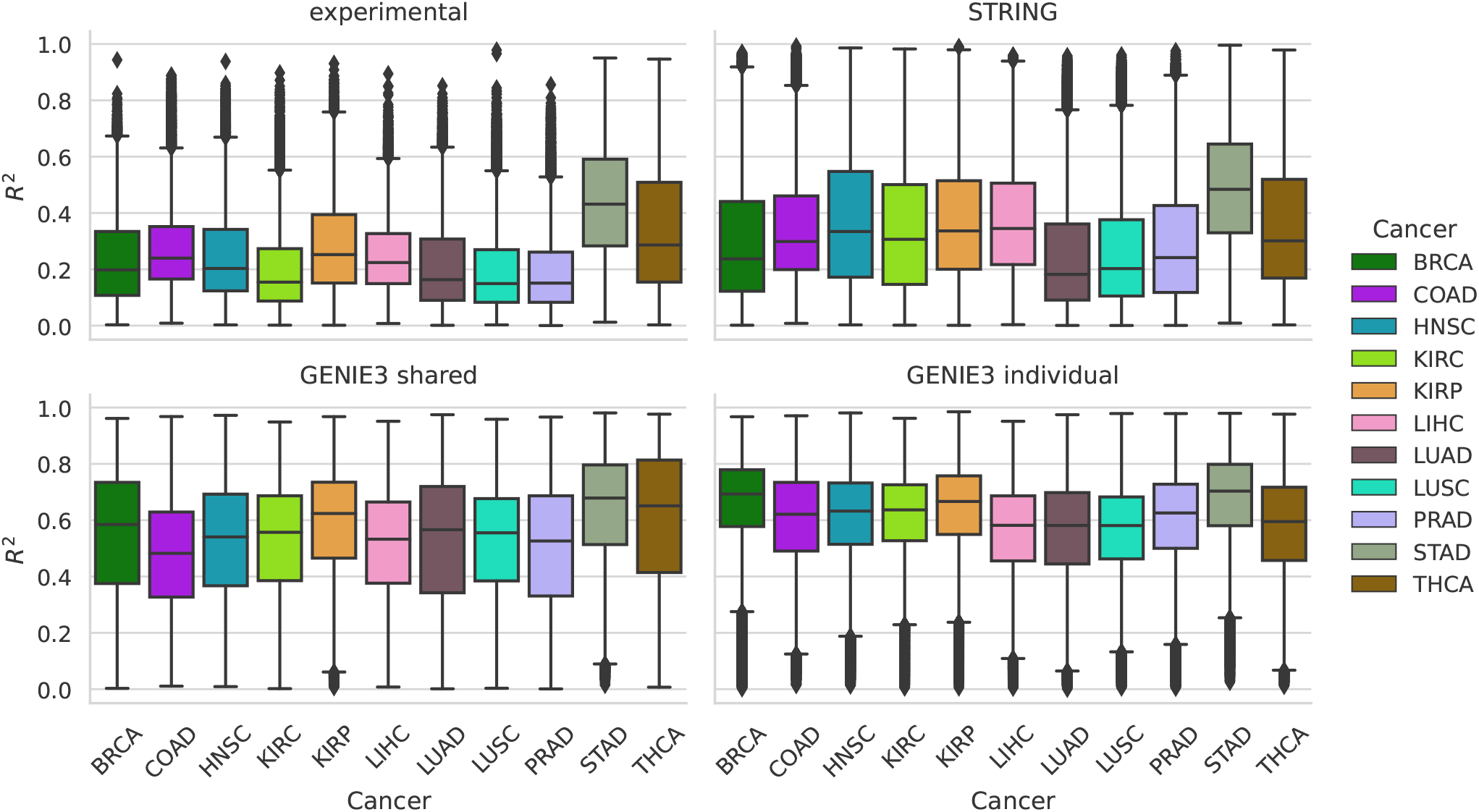
*R*^2^ values representing the goodness of fit of the linear regression models built by DysRegNet for different reference networks. A higher value indicates a better model fit.

**Fig. S3.**
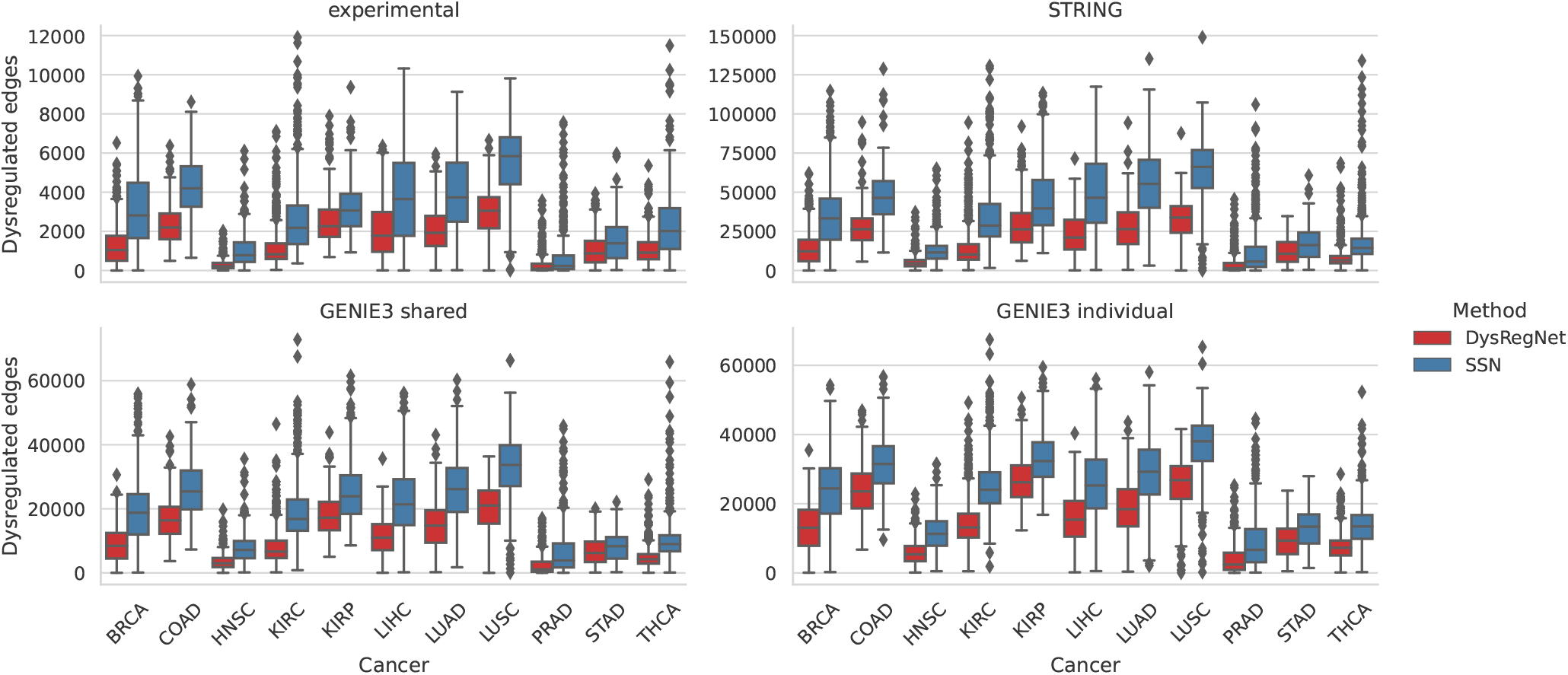
Number of edges in the patient-specific networks inferred with DysRegNet and SSN based on different reference networks.

## Supplementary Note 1: Runtime comparison

For evaluating the time complexity of DysRegNet and SSN, let *n* be the number of control samples, *p* the number of patients/case samples, *g* the number of genes in the expression matrix, *e* the number of edges in a reference network, and *l* the number of covariates used in the linear model of DysRegNet.

SSN computes a correlation matrix for every patient, including the patient and control samples. Using a naive algorithm, calculating the correlation matrix has the computational complexity of 𝒪 (*n* · *g*^2^). Since this procedure has to be repeated for every patient, the total complexity of SSN is 𝒪 (*p* · *n* · *g*^2^). If we do not compute the correlations between every possible gene pair but only those in the reference network, this becomes 𝒪 (*p* · *n* · *e*).

**Fig. S4.**
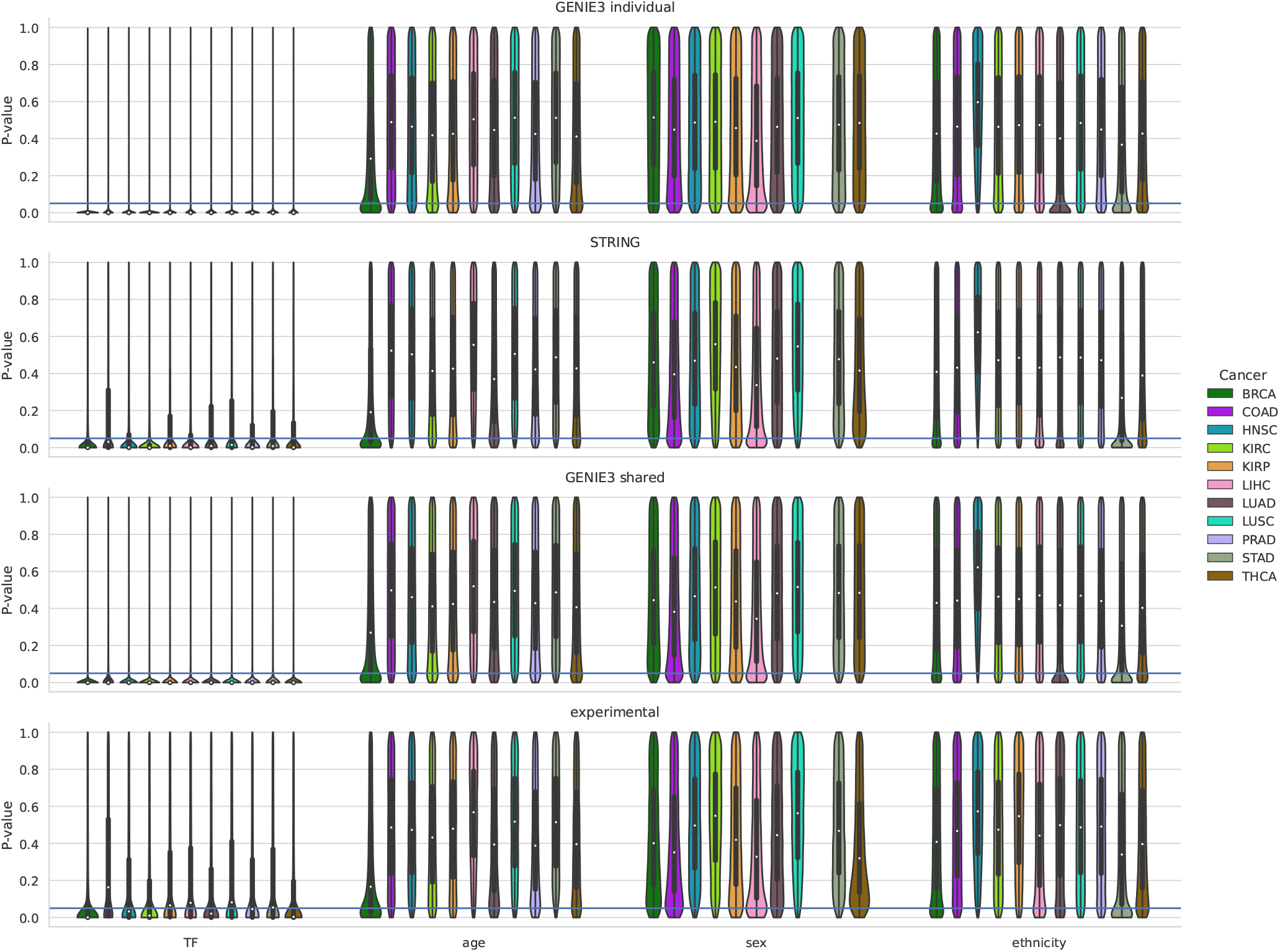
Coefficient p-value distributions obtained with different reference networks (rows) and cancer expression datasets. A single violin summarizes the p-values of a coefficient across all built models (one for each edge in the reference network). The horizontal line indicates a p-value of 0.05.

DysRegNet relies on an ordinary least squares model, the complexity of which depends on the number of control samples *n* and the number of covariates *l*. Using again a naive algorithm, the time complexity of building a single model is 𝒪 ((*n* + *l*) · *l*^2^). Additionally, we have to compute a residual for all patients *p* considering *l* covariates increasing the complexity to 𝒪 ((*n* + *l*) · *l*^2^ +(*p* · *l*)). Suppose we neglect the impact of the number of covariates, which is a reasonable assumption since we would only expect the intercept, the TF expression, and potentially a couple of others. Then, the complexity can be simplified to 𝒪 (*n* + *p*). Building the linear models for every possible gene pair or every edge in the reference network leads to a final complexity of 𝒪 (*g*^2^ · (*n* + *p*)) or 𝒪 (*e* · (*n* + *p*)), respectively.

Comparing the time complexity of DysRegNet and SSN, DysRegNet scales more favorably in terms of the number of control samples and patients, as its effects are additive and not multiplicative, as is the case with SSN.

We also measured the actual runtime of DysRegNet and SSN on the THCA dataset (comprised of 59 control samples and 512 patients) in combination with the experimentally validated HTRIdb [14] reference network. For the benchmark, we kept 9260 genes and 14712 edges by selecting only the genes present in the THCA expression data and the reference network. All methods for the runtime comparison were implemented in Python 3.11, and we measured the total script execution time, including IO operations. Across ten runs, DysRegNet consistently completed in approximately 107 seconds, whereas SSN required about 4500 seconds or 1.25 hours (Figure S6).

**Fig. S5.**
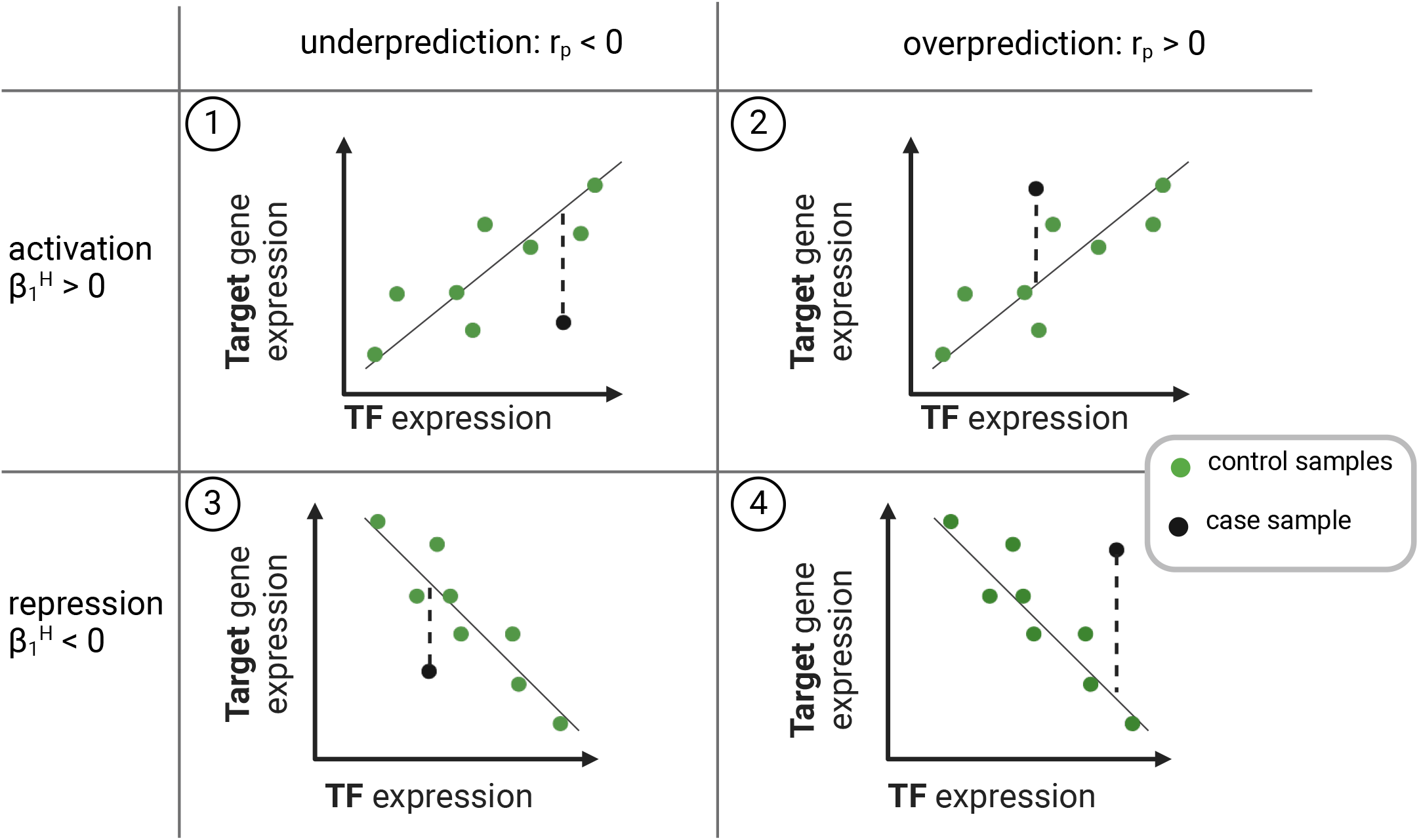
Different scenarios of dysregulation: suppressed activation (1), amplified activation (2), amplified repression (3), and suppressed repression (4).

**Fig. S6.**
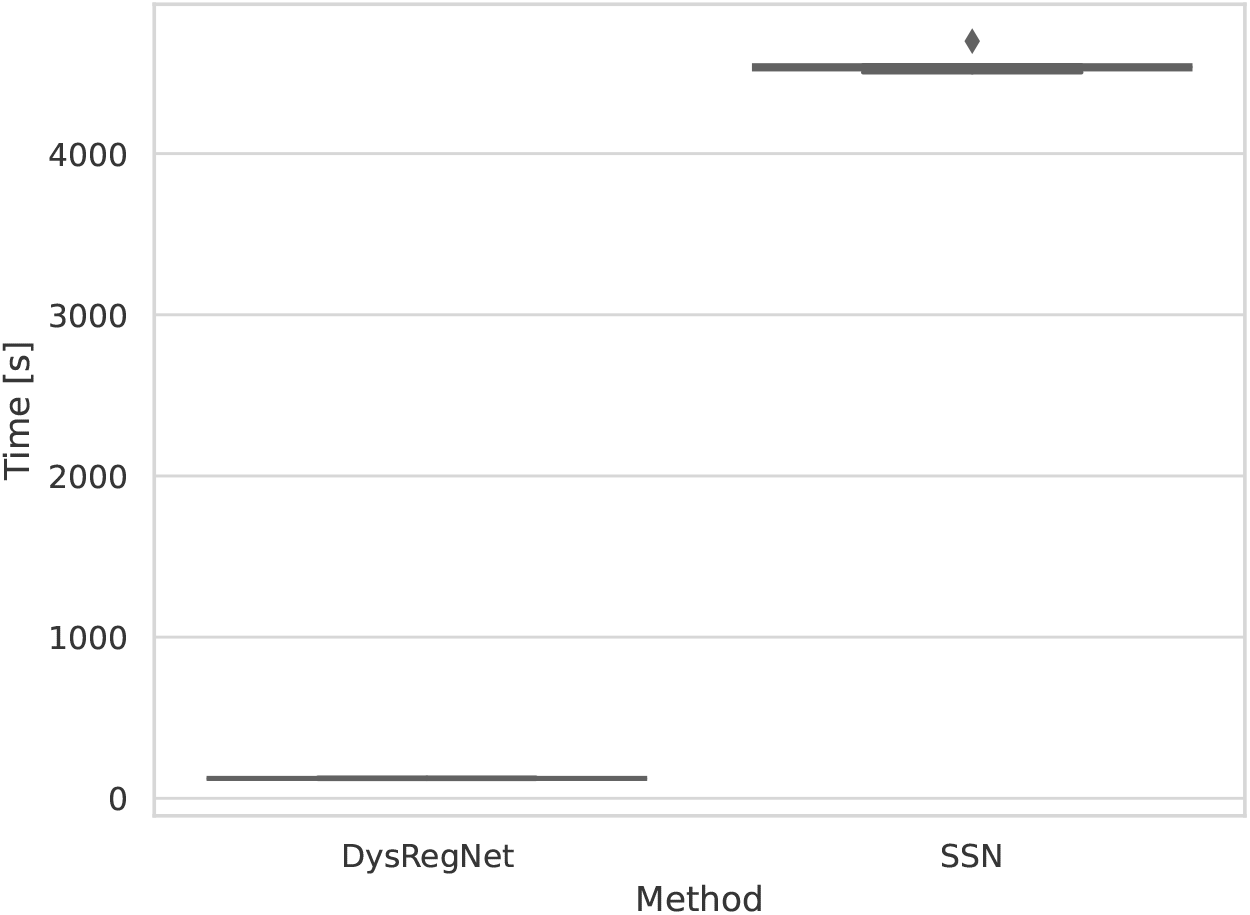
Runtime comparison box plot between DysRegNet and SSN based on ten runs, including 59 control samples, 512 patients, 9260 genes, and 12712 edges in the reference network.

## References

1. Lian En Chai, Swee Kuan Loh, Swee Thing Low, Mohd Saberi Mohamad, Safaai Deris, and Zalmiyah Zakaria. A review on the computational approaches for gene regulatory network construction. Computers in biology and medicine, 48:55–65, 2014.

2. Vân Anh Huynh-Thu, Alexandre Irrthum, Louis Wehenkel, and Pierre Geurts. Inferring regulatory networks from expression data using tree-based methods. PloS one, 5(9):e12776, 2010.

3. Adam A Margolin, Ilya Nemenman, Katia Basso, Chris Wiggins, Gustavo Stolovitzky, Riccardo Dalla Favera, and Andrea Califano. Aracne: an algorithm for the reconstruction of gene regulatory networks in a mammalian cellular context. In BMC bioinformatics, volume 7, pages 1–15. Springer, 2006.

4. Dharmesh D Bhuva, Joseph Cursons, Gordon K Smyth, and Melissa J Davis. Differential coexpression-based detection of conditional relationships in transcriptional data: comparative analysis and application to breast cancer. Genome biology, 20(1):1–21, 2019.

5. Beryl B Cummings, Jamie L Marshall, Taru Tukiainen, Monkol Lek, Sandra Donkervoort, A Reghan Foley, Veronique Bolduc, Leigh B Waddell, Sarah A Sandaradura, Gina L O’Grady, et al. Improving genetic diagnosis in mendelian disease with transcriptome sequencing. Science translational medicine, 9(386):eaal5209, 2017.

6. Laura S Kremer, Daniel M Bader, Christian Mertes, Robert Kopajtich, Garwin Pichler, Arcangela Iuso, Tobias B Haack, Elisabeth Graf, Thomas Schwarzmayr, Caterina Terrile, et al. Genetic diagnosis of mendelian disorders via rna sequencing. Nature communications, 8(1):1–11, 2017.

7. Byungkyu Park, Wook Lee, Inhee Park, and Kyungsook Han. Finding prognostic gene pairs for cancer from patient-specific gene networks. BMC medical genomics, 12(8):1–14, 2019.

8. Wook Lee, De-Shuang Huang, and Kyungsook Han. Constructing cancer patient-specific and group-specific gene networks with multi-omics data. BMC medical genomics, 13(6):1–12, 2020.

9. Marieke Lydia Kuijjer, Matthew George Tung, GuoCheng Yuan, John Quackenbush, and Kimberly Glass. Estimating sample-specific regulatory networks. Iscience, 14:226–240, 2019.

10. Xiaoping Liu, Yuetong Wang, Hongbin Ji, Kazuyuki Aihara, and Luonan Chen. Personalized characterization of diseases using sample-specific networks. Nucleic acids research, 44(22):e164–e164, 2016.

11. Yanhong Huang, Xiao Chang, Yu Zhang, and Xiaoping Chen, Luonan and Liu. Disease characterization using a partial correlation-based sample-specific network. Brief. Bioinform., 22(3), 2021.

12. Hsin-Hua Chen, Chun-Wei Hsueh, Chia-Hwa Lee, Ting-Yi Hao, Tzu-Ying Tu, Lan-Yun Chang, Jih-Chin Lee, and Chun-Yu Lin. SWEET: a single-sample network inference method for deciphering individual features in disease. Brief. Bioinform., 24(2), March 2023.

13. Mai Adachi Nakazawa, Yoshinori Tamada, Yoshihisa Tanaka, Marie Ikeguchi, Kako Higashihara, and Yasushi Okuno. Novel cancer subtyping method based on patient-specific gene regulatory network. Scientific Reports, 11(1):1–11, 2021.

14. Luiz A Bovolenta, Marcio L Acencio, and Ney Lemke. Htridb: an open-access database for experimentally verified human transcriptional regulation interactions. BMC genomics, 13(1):1–10, 2012.

15. Lars J Jensen, Michael Kuhn, Manuel Stark, Samuel Chaffron, Chris Creevey, Jean Muller, Tobias Doerks, Philippe Julien, Alexander Roth, Milan Simonovic, et al. String 8—a global view on proteins and their functional interactions in 630 organisms. Nucleic acids research, 37(suppl_1):D412–D416, 2009.

16. Ezio Laconi, Fabio Marongiu, and James DeGregori. Cancer as a disease of old age: changing mutational and microenvironmental landscapes. British journal of cancer, 122(7):943–952, 2020.

17. Berna C Özdemir and Gian-Paolo Dotto. Racial differences in cancer susceptibility and survival: more than the color of the skin? Trends in cancer, 3(3):181–197, 2017.

18. Hae-In Kim, Hyesol Lim, and Aree Moon. Sex differences in cancer: epidemiology, genetics and therapy. Biomolecules & therapeutics, 26(4):335, 2018.

19. Harold W Kuhn. The hungarian method for the assignment problem. Naval research logistics quarterly, 2(1-2):83–97, 1955.

20. Theresa Phillips et al. The role of methylation in gene expression. Nature Education, 1(1):116, 2008.

21. Emmanouil Bouras, Meropi Karakioulaki, Konstantinos I Bougioukas, Michalis Aivaliotis, Georgios Tzimagiorgis, and Michael Chourdakis. Gene promoter methylation and cancer: An umbrella review. Gene, 710:333–340, 2019.

22. Venkateshwar G Keshamouni. Excavation of fosl1 in the ruins of kras-driven lung cancer, 2018.

23. Xiaodong Sun, Henry F Frierson, Ceshi Chen, Changling Li, Qimei Ran, Kristen B Otto, Brandi M Cantarel, Robert L Vessella, Allen C Gao, John Petros, et al. Frequent somatic mutations of the transcription factor atbf1 in human prostate cancer. Nature genetics, 37(4):407–412, 2005.

24. Motoki Takaku, Sara A Grimm, Bony De Kumar, Brian D Bennett, and Paul A Wade. Cancer-specific mutation of gata3 disrupts the transcriptional regulatory network governed by estrogen receptor alpha, foxa1 and gata3. Nucleic acids research, 48(9):4756–4768, 2020.

25. John H Bushweller. Targeting transcription factors in cancer—from undruggable to reality. Nature Reviews Cancer, 19(11):611–624, 2019.

26. Douglas Hanahan and Robert A Weinberg. Hallmarks of cancer: the next generation. cell, 144(5):646–674, 2011.

27. Douglas Hanahan. Hallmarks of cancer: New dimensions. Cancer Discov., 12(1):31–46, 2022.

28. Chandrika Canugovi, Mark D Stevenson, Aleksandr E Vendrov, Takayuki Hayami, Jacques Robidoux, Han Xiao, You-Yi Zhang, Daniel T Eitzman, Marschall S Runge, and Nageswara R Madamanchi. Increased mitochondrial NADPH oxidase 4 (NOX4) expression in aging is a causative factor in aortic stiffening. Redox Biol, 26:101288, September 2019.

29. Hwa-Young Lee, Hyun-Kyoung Kim, The-Hiep Hoang, Siyoung Yang, Hyung-Ryong Kim, and Han-Jung Chae. The correlation of IRE1α oxidation with nox4 activation in aging-associated vascular dysfunction. Redox Biol, 37:101727, October 2020.

30. Chang Liu, Libangxi Liu, Minghui Yang, Bin Li, Jiarong Yi, Xuezheng Ai, Yang Zhang, Bo Huang, Changqing Li, Chencheng Feng, and Yue Zhou. A positive feedback loop between EZH2 and NOX4 regulates nucleus pulposus cell senescence in age-related intervertebral disc degeneration. Cell Div., 15:2, February 2020.

31. Jenny C Link, Carrie B Wiese, Xuqi Chen, Rozeta Avetisyan, Emilio Ronquillo, Feiyang Ma, Xiuqing Guo, Jie Yao, Matthew Allison, Yii-Der Ida Chen, Jerome I Rotter, Julia S El-Sayed Moustafa, Kerrin S Small, Shigeki Iwase, Matteo Pellegrini, Laurent Vergnes, Arthur P Arnold, and Karen Reue. X chromosome dosage of histone demethylase KDM5C determines sex differences in adiposity. J. Clin. Invest., 130(11):5688–5702, November 2020.

32. Milan Kumar Samanta, Srimonta Gayen, Clair Harris, Emily Maclary, Yumie Murata-Nakamura, Rebecca M Malcore, Robert S Porter, Patricia M Garay, Christina N Vallianatos, Paul B Samollow, Shigeki Iwase, and Sundeep Kalantry. Activation of xist by an evolutionarily conserved function of KDM5C demethylase. Nat. Commun., 13(1):2602, May 2022.

33. Theresa M Filtz, Walter K Vogel, and Mark Leid. Regulation of transcription factor activity by interconnected post-translational modifications. Trends Pharmacol. Sci., 35(2):76–85, February 2014.

34. Sachi Inukai, Kian Hong Kock, and Martha L Bulyk. Transcription factor-DNA binding: beyond binding site motifs. Curr. Opin. Genet. Dev., 43:110–119, April 2017.

35. Ralph B d’Agostino. An omnibus test of normality for moderate and large size samples. Biometrika, 58(2):341–348, 1971.

36. E S Pearson. Tests for departure from normality. empirical results for the distributions of b2 and √ b1. Biometrika, 60(3):613–622, December 1973.

37. Anna Ketteler and David B Blumenthal. Demographic confounders distort inference of gene regulatory and gene co-expression networks in cancer. Brief. Bioinform., 24(6), September 2023.

38. Sarah Kim-Hellmuth, François Aguet, Meritxell Oliva, Manuel Muñoz-Aguirre, Silva Kasela, Valentin Wucher, Stephane E Castel, Andrew R Hamel, Ana Viñuela, Amy L Roberts, Serghei Mangul, Xiaoquan Wen, Gao Wang, Alvaro N Barbeira, Diego Garrido-Martín, Brian B Nadel, Yuxin Zou, Rodrigo Bonazzola, Jie Quan, Andrew Brown, Angel Martinez-Perez, José Manuel Soria, GTEx Consortium, Gad Getz, Emmanouil T Dermitzakis, Kerrin S Small, Matthew Stephens, Hualin S Xi, Hae Kyung Im, Roderic Guigó, Ayellet V Segrè, Barbara E Stranger, Kristin G Ardlie, and Tuuli Lappalainen. Cell type-specific genetic regulation of gene expression across human tissues. Science, 369(6509), September 2020.

39. S Ballouz, W Verleyen, and J Gillis. Guidance for RNA-seq co-expression network construction and analysis: safety in numbers. Bioinformatics, 31(13):2123–2130, July 2015.

40. John Lonsdale, Jeffrey Thomas, Mike Salvatore, Rebecca Phillips, Edmund Lo, Saboor Shad, Richard Hasz, Gary Walters, Fernando Garcia, Nancy Young, et al. The genotype-tissue expression (gtex) project. Nature genetics, 45(6):580–585, 2013.

41. Joke Deschildre, Boris Vandemoortele, Jens Uwe Loers, Katleen De Preter, and Vanessa Vermeirssen. Evaluation of single-sample network inference methods for precision oncology. NPJ Syst Biol Appl, 10(1):18, February 2024.

42. Margherita De Marzio, Kimberly Glass, and Marieke L Kuijjer. Single-sample network modeling on omics data. BMC Biol., 21(1):296, December 2023.

43. Nina Baumgarten, Laura Rumpf, Thorsten Kessler, and Marcel H Schulz. A statistical approach to identify regulatory DNA variations. bioRxiv, February 2023.

44. Héctor Climente-González, Eduard Porta-Pardo, Adam Godzik, and Eduardo Eyras. The functional impact of alternative splicing in cancer. Cell reports, 20(9):2215–2226, 2017.

45. Zakaria Louadi, Kevin Yuan, Alexander Gress, Olga Tsoy, Olga V Kalinina, Jan Baumbach, Tim Kacprowski, and Markus List. Digger: exploring the functional role of alternative splicing in protein interactions. Nucleic acids research, 49(D1):D309–D318, 2021.

46. Theodora Katsila, Georgios A Spyroulias, George P Patrinos, and Minos-Timotheos Matsoukas. Computational approaches in target identification and drug discovery. Comput. Struct. Biotechnol. J., 14:177–184, May 2016.

47. Sepideh Sadegh, Julian Matschinske, David B. Blumenthal, Gihanna Galindez, Tim Kacprowski, Markus List, Reza Nasirigerdeh, Mhaned Oubounyt, Andreas Pichlmair, Tim Daniel Rose, Marisol Salgado-Albarr’an, Julian Sp”ath, Alexey Stukalov, Nina K. Wenke, Kevin Yuan, Josch K. Pauling, and Jan Baumbach. Exploring the sars-cov-2 virus-host-drug interactome for drug repurposing. Nature Communications, 11(1):3518, Jul 2020.

48. Sepideh Sadegh, James Skelton, Elisa Anastasi, Judith Bernett, David B Blumenthal, Gihanna Galindez, Marisol Salgado-Albarrán, Olga Lazareva, Keith Flanagan, Simon Cockell, et al. Network medicine for disease module identification and drug repurposing with the nedrex platform. Nature communications, 12(1):1–12, 2021.

49. Michael Hartung, Elisa Anastasi, Zeinab M Mamdouh, Cristian Nogales, Harald H H W Schmidt, Jan Baumbach, Olga Zolotareva, and Markus List. Cancer driver drug interaction explorer. Nucleic Acids Res., 50(W1):W138–44, May 2022.

50. Cristian Nogales, Zeinab M Mamdouh, Markus List, Christina Kiel, Ana I Casas, and Harald H H W Schmidt. Network pharmacology: curing causal mechanisms instead of treating symptoms. Trends Pharmacol. Sci., 43(2):136–150, February 2022.

51. Mary J Goldman, Brian Craft, Mim Hastie, Kristupas Repečka, Fran McDade, Akhil Kamath, Ayan Banerjee, Yunhai Luo, Dave Rogers, Angela N Brooks, et al. Visualizing and interpreting cancer genomics data via the xena platform. Nature biotechnology, 38(6):675–678, 2020.

52. Samuel A Lambert, Arttu Jolma, Laura F Campitelli, Pratyush K Das, Yimeng Yin, Mihai Albu, Xiaoting Chen, Jussi Taipale, Timothy R Hughes, and Matthew T Weirauch. The human transcription factors. Cell, 172(4):650–665, 2018.

53. F. Pedregosa, G. Varoquaux, A. Gramfort, V. Michel, B. Thirion, O. Grisel, M. Blondel, P. Prettenhofer, R. Weiss, V. Dubourg, J. Vanderplas, A. Passos, D. Cournapeau, M. Brucher, M. Perrot, and E. Duchesnay. Scikit-learn: Machine learning in Python. Journal of Machine Learning Research, 12:2825–2830, 2011.

54. Max Franz, Christian T. Lopes, Gerardo Huck, Yue Dong, Onur Sumer, and Gary D. Bader. Cytoscape.js: a graph theory library for visualisation and analysis. Bioinformatics, 32(2):309–311, 09 2015.

55. Andreas Maier, Michael Hartung, Mark Abovsky, Klaudia Adamowicz, Gary D Bader, Sylvie Baier, David B Blumenthal, Jing Chen, Maria L Elkjaer, Carlos Garcia-Hernandez, Markus Hoffmann, Igor Jurisica, Max Kotlyar, Olga Lazareva, Hagai Levi, Markus List, Sebastian Lobentanzer, Joseph Loscalzo, Noel Malod-Dognin, Quirin Manz, Julian Matschinske, Mhaned Oubounyt, Alexander R Pico, Rudolf T Pillich, Julian M Poschenrieder, Dexter Pratt, Nataša Pržulj, Sepideh Sadegh, Julio Saez-Rodriguez, Suryadipto Sakar, Gideon Shaked, Ron Shamir, Nico Trummer, Ugur Turhan, Ruisheng Wang, Olga Zolotareva, and Jan Baumbach. Drugst. One – a plug-and-play solution for online systems medicine and network-based drug repurposing. May 2023.

